# Evidence for Powassan virus deletions and defective RNA in field collected ticks

**DOI:** 10.1101/2024.09.30.615485

**Authors:** Rose M. Langsjoen, Samantha J. Courtney, Chasity E. Trammell, Rebecca M. Robich, Heidi K. Goethert, Rebekah J. McMinn, Sam R. Telford, Gregory D. Ebel, Anne Piantadosi

**Affiliations:** Department of Pathology and Laboratory Medicine, Emory University, Atlanta, GA; Center for Vector-borne Infectious Diseases, Department of Microbiology, Immunology, and Pathology, Colorado State University, Fort Collins, CO; MaineHealth Institute for Research, Maine Medical Center, Scarborough, ME; Dept of Infectious Disease and Global Health, Tufts University, North Grafton, MA; Division of Infectious Diseases, Department of Medicine, Emory University, Atlanta, GA

**Keywords:** Powassan virus, deer tick virus, tick-borne flavivirus, recombination, deletions, defective virus, defective RNAs, defective viral genomes, structural variants

## Abstract

Powassan virus (POWV) is a tick-borne flavivirus in the tick-borne encephalitis virus (TBEV) serogroup endemic to the United States, Canada, and parts of Russia. POWV remains an under-studied pathogen, despite the potential for serious and life-threatening neurologic complications following infection. While prior studies have characterized viral diversity due to single nucleotide polymorphisms, little is known about POWV recombination, defective RNAs (D-RNAs), and functional structural variants (SVs). Understanding POWV recombination in its natural vector can provide important insights into its replication and evolution. Thus, we analyzed POWV sequence data from 51 ticks collected from the Northeast United States to characterize deletion expression levels and patterns in naturally infected ticks, and we compared these results to single-passage isolates. We found that deletions were common in POWV RNA from ticks and that several areas of the genome were enriched for recombination junctions. Deletions were often associated with areas of microhomology. While most deletions were sample-specific, two major deletion archetypes were observed across multiple tick samples. The first consisted of small 19-50 base deletions in the methyltransferase domain of the ns5 RNA-dependent RNA-polymerase gene, resulting in a mixture of putative SVs and D-RNAs. The second consisted of approximately 1600 base deletions spanning the ns2a-ns3 genes, resulting in putative D-RNAs with abrogated viral protease function. Protease deletions were significantly enriched after one passage in baby hamster kidney cells despite a decrease in overall deletion expression. These results demonstrate the proclivity of POWV for recombination, with potential implications for immune evasion and persistence in ticks.

**IMPORTANCE:** Powassan virus is a tick-borne flavivirus that can cause serious, life-threatening neurological disease. Understanding how Powassan virus replicates and evolves within its tick vector may elucidate factors important in persistence, transmission, and human disease. Defective RNAs are replication-incompetent viral genomes generated through internal deletions, which have been associated with disease severity and persistent infection in other viruses but have not been described for Powassan virus. Here, we show that Powassan virus produces abundant defective RNAs in field-caught ticks, and that expression patterns of these defective RNAs changes after one passage in mammalian cells. Although the function of these defective RNAs remains unknown, this work establishes a critical framework for investigating the role of defective RNAs in Powassan virus replication and transmission.

## Introduction

Powassan virus (POWV) is a flavivirus within the tick-borne encephalitis virus (TBEV) serogroup endemic in the US, Canada, and the Russian Far East [1–4]. POWV is an emerging pathogen of concern [5], given the increase in disease incidence over the past decade and the ability of the virus to cause severe and potentially fatal viral encephalitis. While comparative genomics of POWV isolates has yielded critical data for understanding POWV evolution, few studies have considered the diversity of POWV populations within individual hosts, and still fewer have considered the recombinant components of those populations. Viral RNA recombination can have a profound influence on viral evolution, as well as produce defective virus that can impact viral replication, persistence, and disease severity. Thus, by characterizing viral recombination in relevant POWV hosts, we can determine potential sources of genetic variation and identify novel research avenues for identifying determinants of persistence and virulence.

Recombination is a key driver of RNA virus evolution [6], including for mosquito-borne flaviviruses [7, 8]. This process involves template switching between distinct RNA strands or within the same strand and can generate deletions and duplications [9] that result in RNA structural variants (SVs) or defective RNAs (D-RNAs; also known as defective viral genomes, DVGs, or defective-interfering RNAs, DI-RNAs) [10, 11]. Both SVs and D-RNAs are recombinant viral RNAs with deletions and/or duplications, but they differ functionally: SVs remain autonomously functional, while D-RNAs cannot replicate and/or transmit autonomously [12–14]. Although SVs and D-RNAs were initially thought to be artifacts of *in vitro* passaging, subsequent studies using deep sequencing detected and characterized them in animal models of mosquito-borne flaviviruses and alphaviruses [15–17] and natural infection of respiratory viruses [18–20] and flaviviruses [11]. D-RNAs in tissue culture can have a profound effect by disrupting viral replication and inducing host pro-inflammatory responses [21]. However, in natural infection, D-RNAs may have variable roles in pathogenesis [21], with some studies suggesting therapeutic potential for SARS-CoV-2 and Zika virus [16, 22], others demonstrating a role in promoting persistent infection for Usutu and Langat viruses [15, 23], and still others suggesting differential roles for transmitted versus *de novo* produced alphavirus D-RNAs [17]. Despite the significant global health impact of tick-borne flaviviruses and the potential importance of ticks as reservoirs, our understanding of recombination in these viruses remains limited [24–26], especially the formation of D-RNAs in natural tick vectors.

*Ixodes* ticks vector POWV via bloodmeal exchange with mammalian hosts (e.g. shrews, deer mice, groundhogs, squirrels) and humans [1]. Relatively little is known about the maintenance of POWV and other tick-borne flaviviruses in mammalian reservoirs, and evidence suggests that tick-borne flaviviruses can also be effectively maintained in tick populations [27, 28]. Ticks are unique among arthropod vectors due to their multi-stage, blood-meal dependent life cycle: between each of three active life stages (larva, nymph, and adult), ticks take a single bloodmeal from a single host before molting into the next stage. POWV has been identified in field-collected ticks of multiple life stages [1, 29], and POWV have been shown to traverse life stages through transstadial [30–32] and even transovarial transmission [28, 33]. This is accomplished necessarily through persistent infection of the tick, which can last hundreds of days in laboratory-infected ticks [30, 34]. Because the tick vector may act as an important reservoir for POWV, it is a critical host in which to study POWV replication and recombination.

This study seeks to expand our knowledge of POWV recombination in naturally infected ticks, towards the goal of gaining insight into viral pathogenesis and evolution. We performed detailed recombination analyses to: 1) characterize POWV deletions in naturally infected ticks; 2) determine potential factors that influence recombination leading to deletions; and 3) determine whether tissue culture passage affects POWV deletion content.

## Results

### Deletions occur throughout the POWV genome and most are predicted to result in defective RNAs

We analyzed POWV reads generated by RNA metagenomic short-read sequencing from 53 individual POWV-positive ticks collected from the Northeast U.S. between 2018 and 2020 [29] (**Table 1**; sequencing metrics **Table S1**). Most of the samples (n=33) were collected from Maine in 2018 (n=3), 2019 (n=14), and 2020 (n=16). The remaining samples were collected in either Massachusetts (2018, n=2; 2020, n=6), New York state (2019, n=9), or New Jersey (2019, n=3).

**Table 1.**
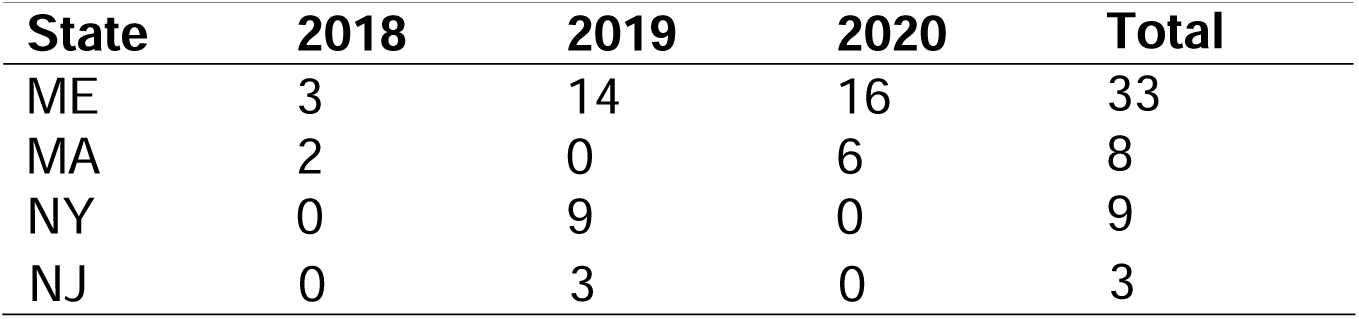
Naturally infected ticks included, by year and origin.

We identified 330 unique deletions across the POWV genome in all samples (**Fig. 1**). Most of these were specific to a single sample, which ranged in expression from 2 to 480 counts/10^6^ mapped reads (**Fig. 1A**); this is suggests *de novo* production on a per-tick basis rather than accumulation over multiple transmission cycles between vector and host species, as has been suggested for some mosquito-borne viruses [35]. On a gene level, there was a large concentration of deletions within the ns5 gene, which is expected given its size relative to the rest of the genome (**Fig. 1B**). There were also a large number of deletions with a 5’ recombination breakpoint junction (Junction 1, J1) in the envelope (E) gene, as well as deletions with a 3’ recombination breakpoint junction (Junction 2, J2) in the ns3 gene (**Fig. 1B**), suggesting that there could be junction enrichment in specific genomic regions. Some specific deletions were observed across multiple samples, the most common which included: deletions in the methyltransferase (MTase) region of the ns5 RNA-dependent RNA-polymerase (RdRp) gene; deletions spanning the viral protease (ns2a-ns3); small deletions in ns1; and deletions spanning the majority of the non-structural region between ns2A and ns5 (**Fig. 1A**). Interestingly, a common deletion archetype previously described for flaviviruses—deletions spanning the envelope (E) gene and ending at the beginning of the ns1 gene [11, 36, 37]—were rare among these samples.

**Figure 1.**
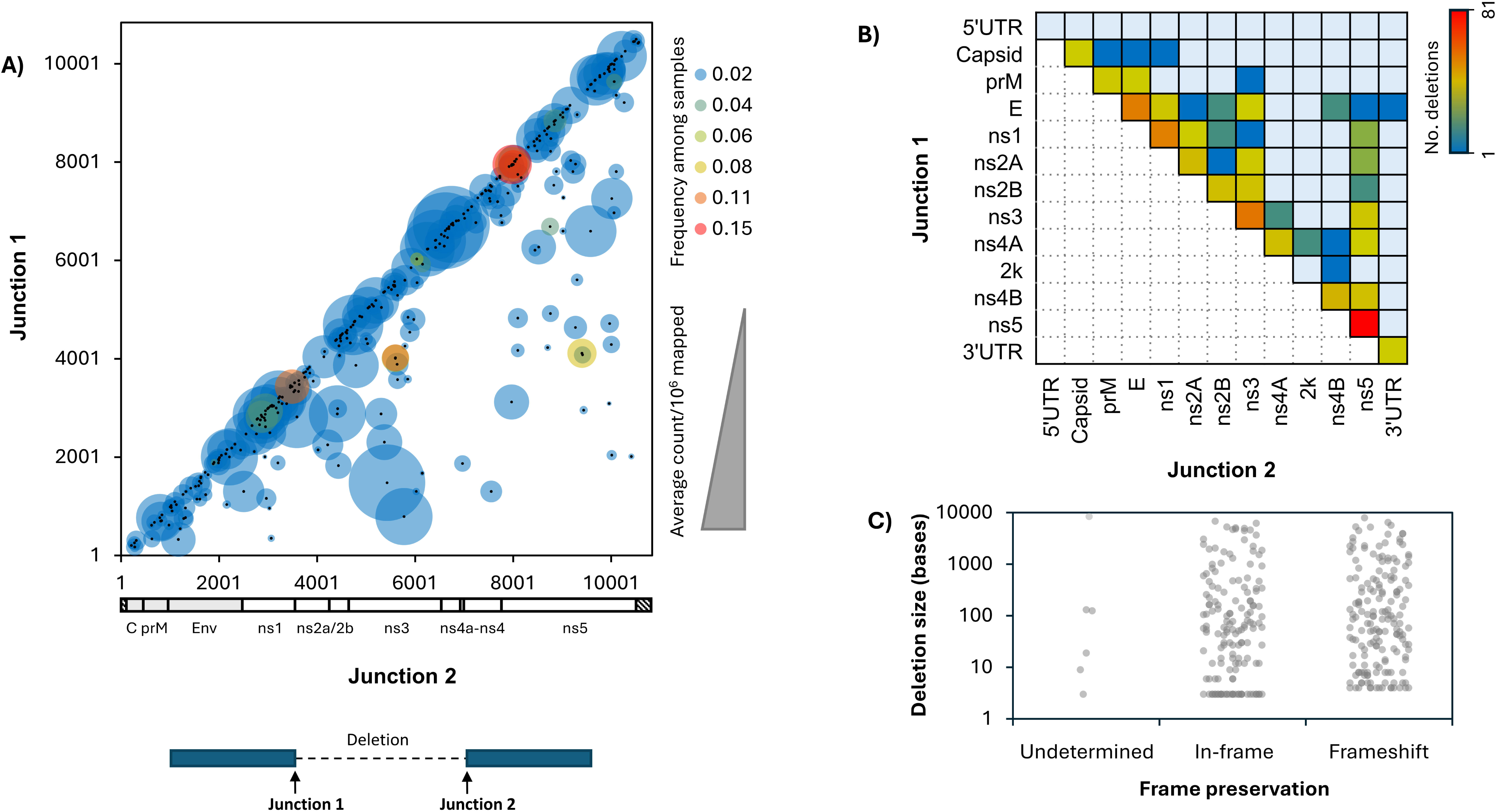
Characteristics of recombination breakpoint junctions and deletions. POWV RNA derived from 53 naturally infected tick samples was sequenced on an Illumina platform, and recombination events were mapped using Viral Recombination Mapper (ViReMa). Deletions were extracted, and recombination breakpoint junction nucleotide positions (Junction 1, Junction 2) were indexed to the POWV reference HM440559.1. **A)** Nucleotide positions of Junction 1 (y-axis) and Junction 2 (X-axis) for each unique deletion were plotted and colored by frequency among samples and sized by average count per million mapped reads; **B)** Total number of deletions by genomic location of Junction 1 (Y-axis) and Junction 2 (X-axis); **C)** Distribution of deletion size (Y-axis) by coding frame (**undetermined**, deletions occurring between non-coding and coding regions of the genome; **in-frame**, deletions with a base-size divisible by 3; **frameshift**, deletions predicted to result in a shift to the coding frame).

Deletions can result in autonomously functional structural variants (SVs) or defective RNAs (D-RNAs; also defective viral genomes, DVGs), which require helper virus to replicate and/or transmit. To evaluate whether the deletions we observed might represent SVs or D-RNAs, we considered their size and coding frame preservation (**Fig. 1C**). Over half of all deletions found in the POWV single open reading frame (ORF) resulted in a frameshift, suggesting they would result in D-RNAs (**Fig. 1C**). A further 23% of all deletions were over 500 nucleotides in length (**Fig. 1C**), and would be expected to disrupt necessary viral proteins, also resulting in D-RNAs. On the other hand, 10% of all deletions observed were only 3 nucleotides in length, resulting in a single amino acid deletion (**Fig. 1C**), which would largely be predicted to result in SVs.

### Common deletion archetypes occur in the ns2A-ns3 protease and methyltransferase domain of the ns5 RNA-dependent RNA-polymerase

Comparing deletion positions between samples, we identified several POWV recombination archetypes that were present in multiple naturally infected ticks (**Fig. 2**). Two of the most common deletions shared similar breakpoint junctions, with the first (5’) in the ns2A gene and the second (3’) in the ns3 gene (**Fig. 2 top,** inset I). These two deletions were collectively present in almost 20% of all tick samples and also corresponded to a region with high overall deletion frequency (**Fig. 2 bottom,** inset I). These deletions are expected to result in the partial removal of the ns3 viral protease as well as the complete removal of its viral cofactors ns2A and ns2B, and are therefore predicted to function as D-RNAs.

**Figure 2.**
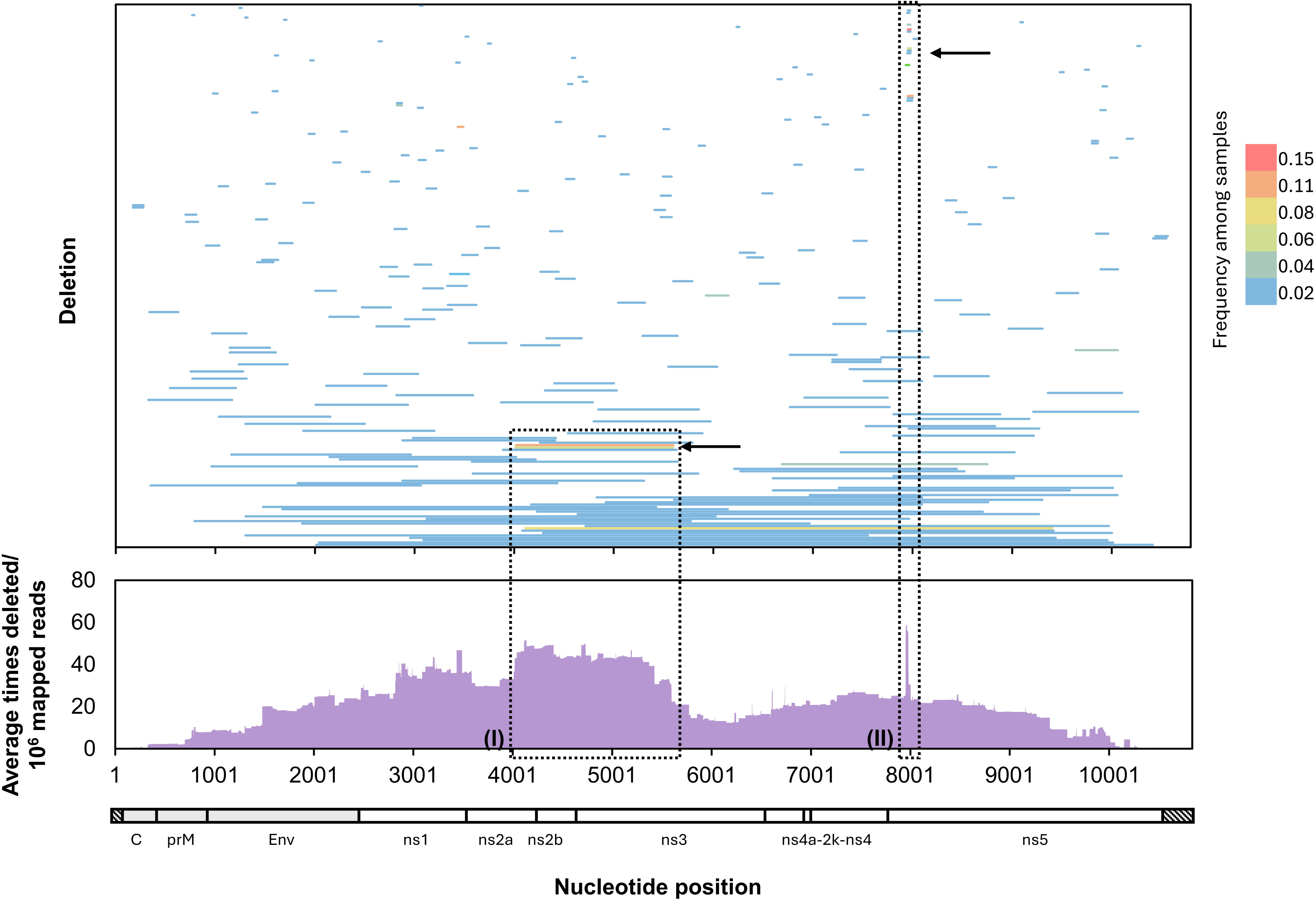
POWV deletion archetypes in naturally infected ticks correspond to peaks in cumulative nucleotide deletion. Individual deletions identified in naturally infected ticks were mapped and colored by frequency among 51 samples (**top**). For each nucleotide position, the number of times it was deleted in each sample was calculated and averaged across all 53 samples (**bottom**). **Inset (I)** highlights a peak in nucleotide deletion frequency across the ns2a-ns3 genes, and **inset (II)** highlights a peak in nucleotide deletion frequency driven by small deletions in the methyltransferase region of the ns5 gene.

The most common single deletion, which was identified in 15% of all samples, occurred between nucleotides 7953 and 7981 in the methyltransferase (MTase) region of the ns5 gene (**Fig. 2 top,** inset II). In addition, eleven other deletions with similar boundaries occurred in the same region. Altogether, MTase deletions between 19 and 50 bases in length between nucleotide positions 7932 and 8004 were identified in over 30% of samples (**Fig. 2 top**). These MTase deletions corresponded to a peak in overall deletion frequency across all samples (**Fig. 2 bottom,** inset II). Interestingly, while the most common MTase deletion is predicted to result in an SV with an in-frame 9 amino acid deletion, most of the others result in RdRp frameshifts and therefore D-RNAs. The deletions occur in a region of the MTase domain that is believed to interact with the RNA-dependent polymerase (RDP) domain in a predicted model of the POWV RdRp (**Fig. 3**). Deleting amino acids at the MTase-RDP interface in Japanese encephalitis virus (JEV) results in significantly decreased, though intact, replication [38], so it is possible that in-frame deletions here would have a similar impact.

**Figure 3.**
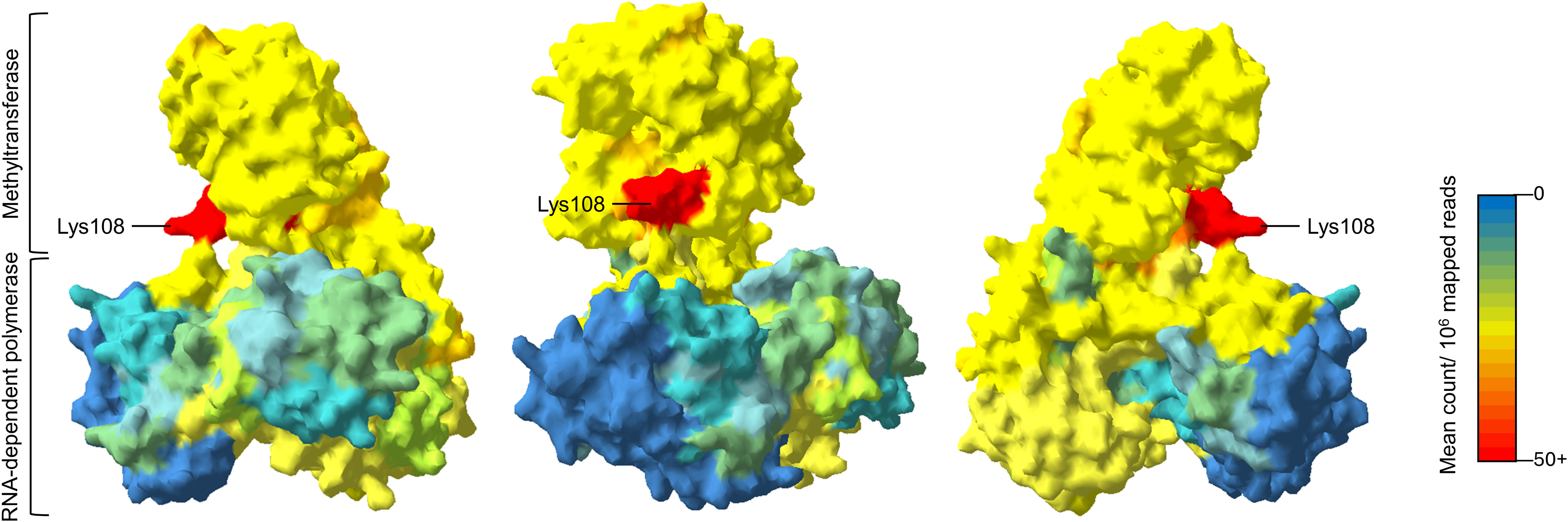
Frequency of amino acid deletion within the POWV ns5 RNA-Dependent RNA-polymerase (RdRp). The POWV ns5 RdRp protein was modeled based on the complete PDB structure for the Japanese encephalitis virus RdRp (PDB 4MTP; Lu G and Gong P 2013 *PLoS Path*) using the ExPasy/SWISS-MODEL tool (Schwede et al 2003 Nucleic Acids Res). Each amino acid was colored by the average number of times it was deleted across all 51 naturally infected ticks. Lysine108 is indicated as a reference point.

### POWV deletions occur at sites of microhomologies between nucleotide junctions

To assess the role of nucleotide sequence in POWV recombination, we evaluated the nucleotide composition and sequence identity at recombination junctions for deletions ≥100 bases in length (**Fig. 4**). Smaller deletions were excluded in order to avoid biases from differential mechanisms regulating small, local deletions such as polymerase slippage. While a slight enrichment of Us was observed immediately down-stream of J1 (**Fig. 4A, top**) and a slight enrichment of Gs was observed upstream of both junctions (**Fig. 4A, bottom**), no specific nucleotide sequence or motif was associated with either junction. However, we found evidence for high nucleotide sequence identity between the 5 nucleotides upstream of J1 and the 5 nucleotides upstream of J2 in most deletions (**Fig. 4B**, **Fig. 4C**). This suggests that microhomologies between breakpoint junctions play a role in template selection during POWV RNA recombination for many but not all deletions.

**Figure 4.**
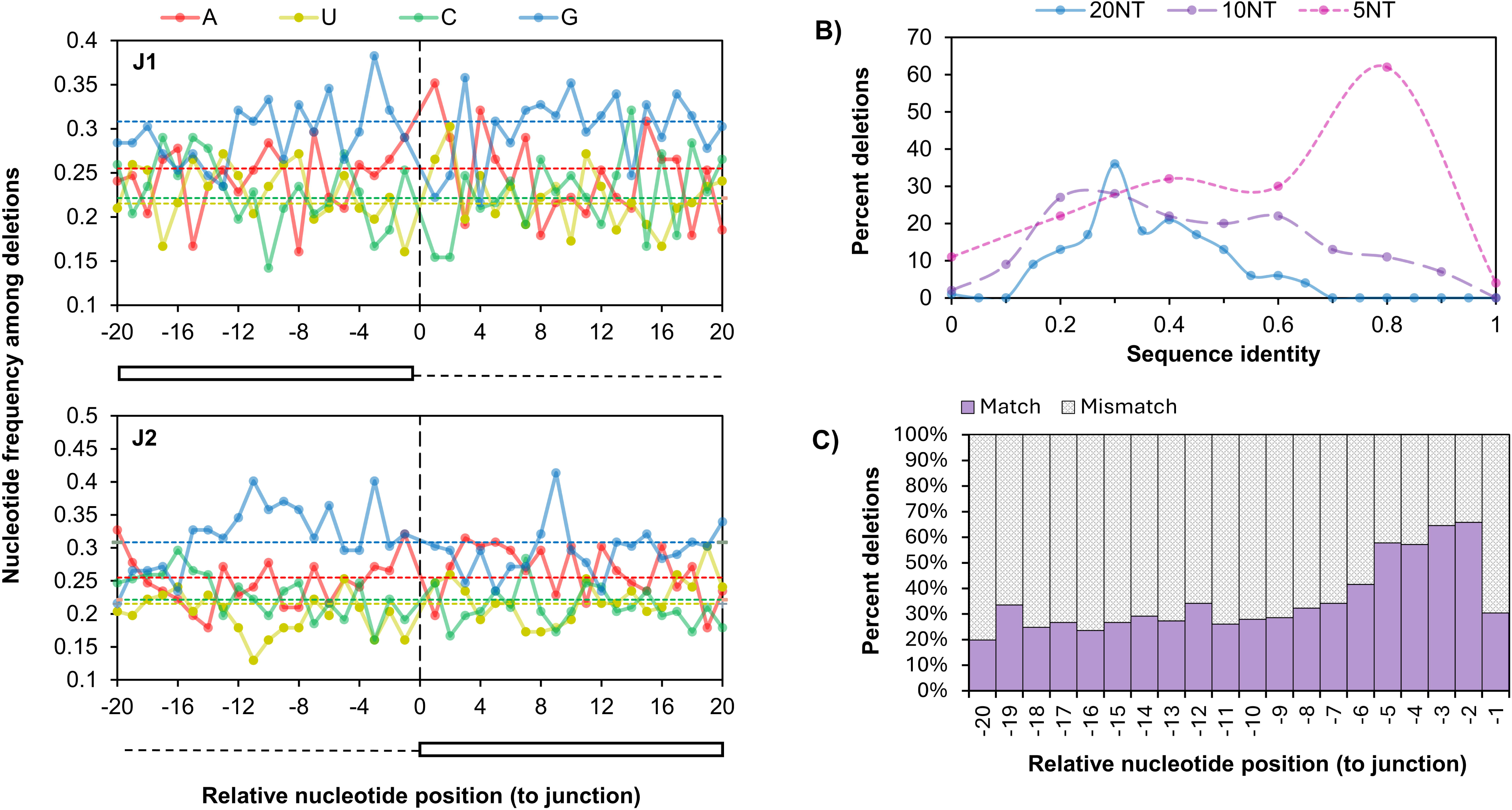
Sequence characteristics at recombination breakpoint junctions identified in naturally infected POWV ticks. **A)** The distribution of bases (solid lines) within 20 bases up- and -downstream of recombination junctions 1 (J1; top) and 2 (J2; bottom) compared to the relative distribution of each base across the POWV genome (dashed lines). **B)** Percent of deletions (Y-axis) with nucleotide sequence identities (X-axis) between junction 1 (the sequence observed upstream of J1 present in the read) and J2 (observed upstream of junction 2 present in the reference) within 5 (pink), 10 (purple), and 20 (blue) nucleotides of the recombination junction. **C)** Percent of deletions with a match (purple) or mismatch (grey) at each nucleotide position relative to the recombination junction.

### POWV protease deletions are significantly enriched after one passage in tissue culture

Because tissue culture passaging has been reported to increase D-RNAs in other viruses, we compared deletions between naturally infected ticks and culture supernatants after one passage in baby hamster kidney (BHK) cells for 13 samples [29]. We did not observe a statistically significant difference in the number of unique deletions (Wilcoxon rank test, p=0.064; **Fig. 5A**) or the total number of deletions (Wilcoxon rank test, p=0.11; **Fig. 5B**), though both tended to decrease after a single passage. Although there was no difference in MTase deletion expression (Wilcoxon rank test, p=0.6; **Fig. 5D**), there was a trend towards higher expression of protease deletions after passage (Paired-Sample Sign Test, p=0.077, **Fig. 5E**). Sequencing depth was not significantly correlated with normalized deletion counts (Kendall test, tau=0.006, p=0.96; **Fig. 5F**), so we do not expect this to be a confounding variable.

**Figure 5.**
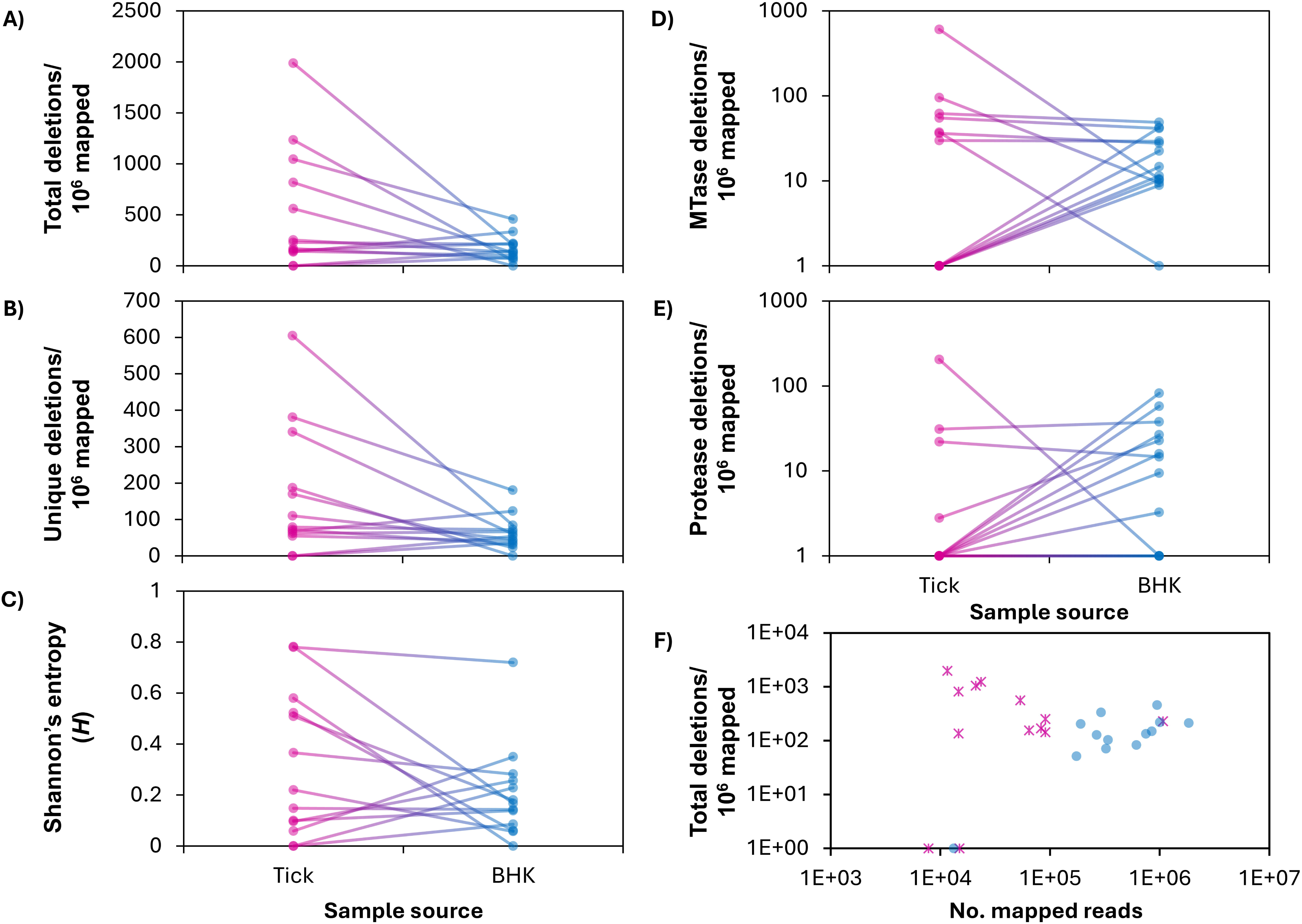
Comparison of Powassan virus deletions between naturally infected ticks and single passage tissue culture supernatant. POWV RNA from naturally infected ticks (**tick**) and RNA derived from supernatant after one passage in baby hamster kidney cells (**BHK**) was sequenced for 13 POWV isolates, and then deletion content was determined by recombination mapping. **A)** Total number of deletions per million mapped reads; **B)** Number of unique deletion species per million mapped reads; **C)** Diversity of deletion species, as determined by Shanon’s entropy (*H*); **D)** Total number of small deletions between 19 and 50 bases in length between nucleotide positions 7932 and 8004 (methyltransferase, MTase deletions) per million mapped reads; **E)** Total number of approximately 1600 base deletions occurring between the ns2a and ns3 genes (protease deletions) per million mapped reads. The total number of deletions per million mapped reads was also compared to the total number of reads per sample **(F**), and the two were not significantly correlated (Kendall test, tau=0.006, p=0.96)

To determine if specific deletions were differentially expressed between naturally infected ticks and single-passage isolates, differential gene (deletion) expression analysis was performed in DESeq2 (**Fig. 6**). Principal component analysis (PCA) revealed that 12/13 tick samples clustered closely together based on deletion expression, while 5/13 BHK samples clustered closely with tick samples and the remaining 8/13 BHK samples showed little to no clustering pattern (**Fig. 6A**). Additionally, very few tick/BHK sample pairs clustered closely with one another. This suggests that, when excluding deletions specific to a single sample, tick samples are more similar to each other than they are to BHK samples. Hierarchical clustering analysis supported this result, with two naturally infected tick sample groups clustering closely together and multiple BHK groups clustering more distantly (**Fig. 6B**). Finally, when evaluating differential expression of specific deletions, two deletions had significantly higher expression in BHK cells: an in-frame 1578 base deletion between nucleotides 4020 and 5599 (log2fold change=-1.5, padj=0.03; **Fig. 6B** inset *, **Fig. 6C** inset I) and a 1582 base frameshift deletion between nucleotides 4020 and 5603 (log2fold change=-1.6, padj=0.04; **Fig. 6B** inset *, **Fig. 6C** inset I). Both are protease deletions and are predicted to function as D-RNAs. Three other deletions—two MTase deletions (**Fig. 6B** inset #, **Fig. 6C** inset II) and one protease deletion (**Fig. 6B** inset #, **Fig. 6C** inset I)—were more highly expressed in BHK cells, though this difference was not statistically significant (−1<log2foldchange<-1.4; 0.05<p<0.38; **Fig. 6C**). No specific deletion species were significantly more highly expressed in tick samples than BHK samples.

**Figure 6.**
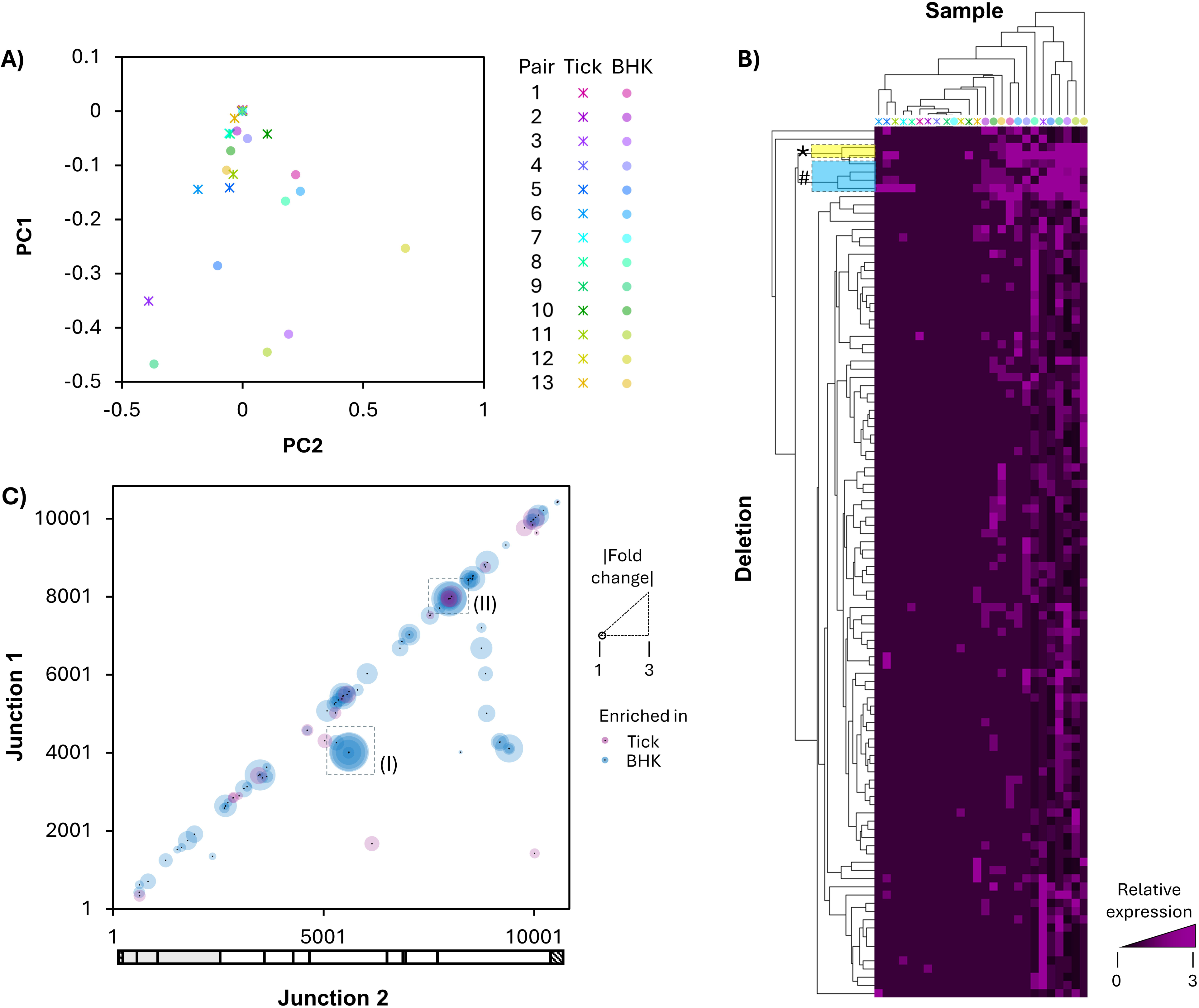
Differential expression of POWV deletions between naturally infected ticks and after one passage in tissue culture. POWV RNA from naturally infected ticks (**tick**) and RNA derived from supernatant after one passage in baby hamster kidney cells (**BHK**) was sequenced for 13 POWV isolates, and then deletion content was determined by recombination mapping. Differential gene expression of individual deletion species was evaluated using DESeq2 software. **A)** Principal component analysis (PCA) was performed, and eigenvalues for components 1 (Y-axis) and 2 X-axis) were plotted for both tick (stars) and BHK (circles) samples and colored by tick/BHK sample pair. **B)** Hierarchical clustering was performed with clustering by sample (X-axis) and deletion (Y-axis). **C)** Each deletion species included in DESeq2 analysis was plotted by recombination junction (Junction 1, Y-axis; Junction 2, X-axis), colored by enrichment by compartment (pink, tick; blue, BHK), and sized by log2 fold change; inset (I) highlights protease deletions, while inset (II) highlights MTase deletions.

### ClickSeq confirms ns2A-ns3 deletions in POWV+ tick samples

To confirm the results of the above analyses, which used previously-generated RNA metagenomic sequencing data, we made additional libraries from samples with available RNA (n=16) using ClickSeq, a library preparation method specifically designed to reduce artifactual recombination events [39, 40]. Analysis of ClickSeq libraries confirmed the presence of ns2A-ns3 protease deletions and identified more unique species of protease deletions and higher levels of expression (**Fig. 7A-B**; insets A.I and B.I). We were not able to confirm the presence of MTase deletions because RNA was not available from the samples with high levels of MTase deletion expression in previous metagenomic libraries: only 4 of the 16 samples that were analyzed using ClickSeq had MTase deletions at low levels (**Fig. 7**).

**Figure 7.**
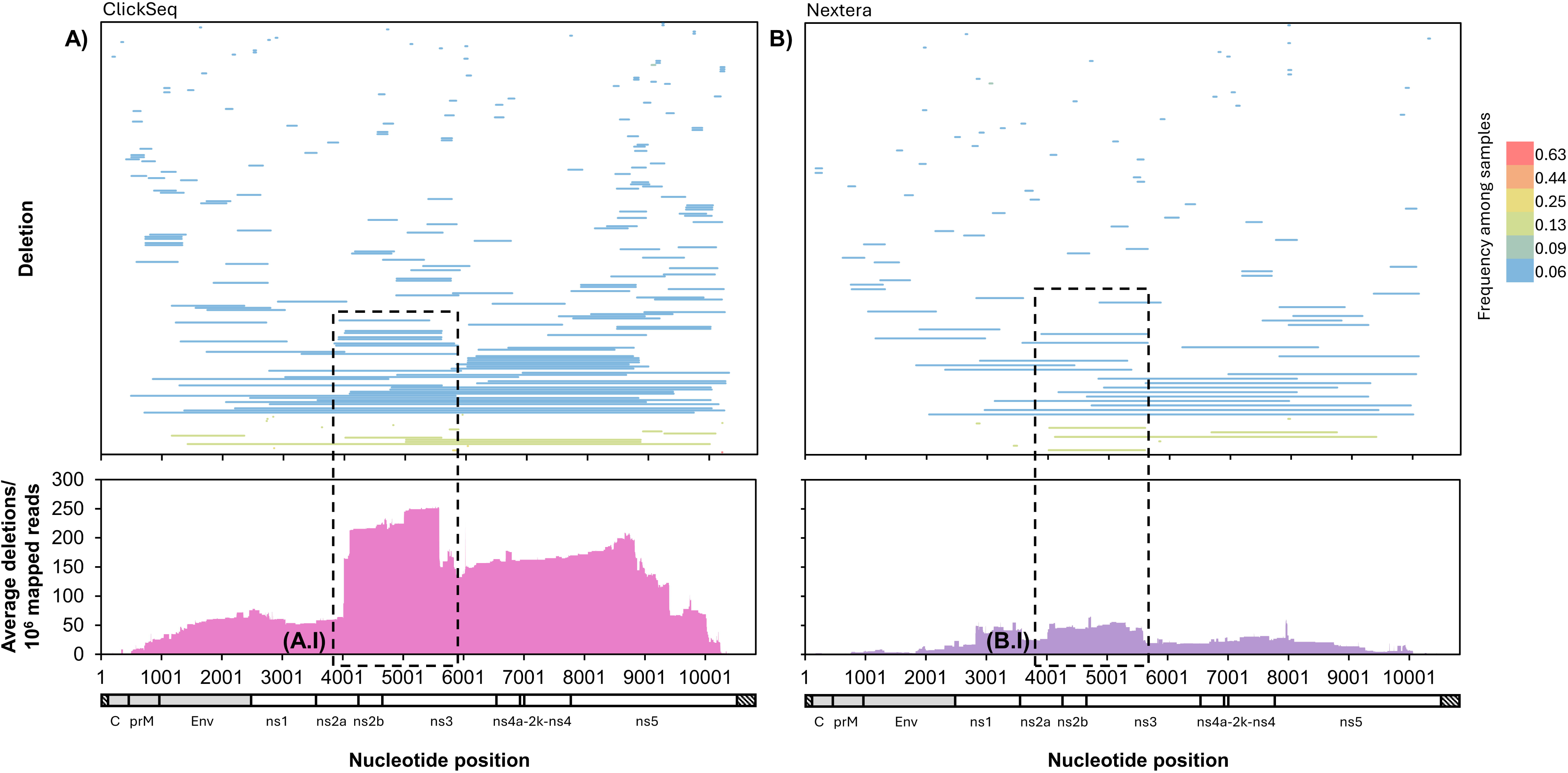
Comparison of POWV deletion expression using two different library preparation methods. POWV RNA from 16 naturally infected tick samples was sequenced using ClickSeq (**A**) and a previous metagenomic approach (**B**). Individual deletion species were mapped by genome position and colored by frequency across samples (**top**). For each nucleotide position, the number of times it was deleted in each sample was calculated and averaged across all 16 samples (**bottom**).

## Discussion

Deletions are common in RNA viruses and are thought to be the result of copy-choice non-homologous RNA recombination; they can result in defective RNAs (D-RNAs), dysfunctional RNAs that can affect wild-type virus replication, as well as autonomously functional structural variants (SVs). Despite the importance of characterizing deletions in RNA viruses, little work has been done to characterize intrahost recombinant RNAs of tick-borne flaviviruses, especially in the natural tick vector. Here, we found that the tick-borne flavivirus Powassan virus (POWV) produces frequent deletions with a range of sizes throughout the genome, leading to both putative D-RNAs and SVs. We found evidence for small 2-5 base homologies between recombination junctions, suggesting that microhomologies may impact template selection during recombination. Importantly, we found several deletion archetypes that were present in multiple samples, including deletions in the ns2-ns3 regions spanning the viral protease and its cofactors, as well as small deletions in the methyltransferase (MTase) region of the ns5 gene. Deletion expression did not increase after a single passage in tissue culture and rather tended to decrease, except for protease deletions, which tended to increase after a single tissue culture passage. Finally, we validated the presence of deletions using a recombination-specific sequencing method. In all, we provide compelling evidence that POWV produces SVs and D-RNAs in a *de novo* fashion during replication in its natural tick vector, and that some deletions arise repeatedly across multiple samples.

While RNA virus recombination and deletions have been described in mammalian hosts, little work has been done to characterize recombinant populations in arthropods; still fewer studies have considered recombinant RNAs in tick hosts. Phylogenetic studies of tick-borne encephalitis virus (TBEV) and louping ill virus (LIV), both close relatives of POWV, have indicated potential recombination events in the evolutionary history of these viruses [24–26, 41, 42], although it is not known in which host they may have occurred. Mosquito-borne flaviviruses have shown differential proclivities for RNA recombination and deletion production both *in vitro* and *in vivo*. Prominently, D-RNAs consisting of deletions between the prM and ns1 genes have been described for West Nile Virus (WNV) in lorikeets [11] and Japanese encephalitis virus [37] and Murray Valley fever virus [36] in tissue culture, where they have been associated with inhibition of wild-type virus replication [11] and establishment of persistent infections [36, 37]. Although the function of the POWV deletions identified in this study remains unclear, our identification of deletions among POWV populations highlights the recombinatorial activity during tick-borne flavivirus replication and recommends further exploration of the role of recombination in tick-borne flavivirus evolution and pathogenesis.

We also show that POWV deletion archetypes arise independently in tick samples from different geographic locations collected at different times. Prominent among these were deletions between the ns2A and ns3 regions, which would delete the viral protease and its cofactors and thus be predicted to function as D-RNAs. Our results additionally show that expression of protease deletions significantly increases after one passage in mammalian cells. It is thus interesting to consider whether these POWV protease deletions may play a host-specific role during POWV replication or be by-product of processes critical to replication, such as RNA-RNA and/or protein-RNA interactions. For example, previous studies have shown that the tick-borne flavivirus protease triggers cellular antiviral responses through its interaction with human TRIM-5α for TBEV and Langat virus (LGTV), but not POWV [43]. While specific TRIM-family proteins and their functions are highly host- and virus-specific [44], it is possible that deleting this region may aid POWV in evading mammalian TRIM-mediated immunity. This recommends the comparison of D-RNA populations between TBEV, LGTV, and POWV to determine potential roles for protease deletions during viral replication.

While POWV protease deletions increased in expression after one passage in tissue culture, overall deletion content was lower in single passage isolates than ticks. This is an intriguing finding, as the current paradigm is that D-RNAs accumulate over high-multiplicity of infection (MOI) tissue culture passage. It is possible that a low MOI enriched the wild-type population in this case, but the MOI is unknown as whole tick homogenate was used for isolation in the absence of quantitative assessment of infectious titer. There are also several potential hypotheses for this result. The first is that tick homogenates include both intracellular and extracellular viral population, and the decrease in cell culture deletion content is driven by a lack of non-packaged virus in supernatant. Previous work with influenza virus [45] and multiple alphavirus species [17] has shown that intracellular viral populations are more recombination-rich than extracellular populations in low-passage tissue culture experiments, as well as in a mouse-model of chikungunya virus infection [17]. Recombinant POWV populations may likewise be enriched intracellularly; this hypothesis can be explicitly tested experimentally by comparing intracellular and extracellular recombinant POWV RNA populations in similarly designed passaging studies or in carefully designed *in vivo* models of infection.

The second potential hypothesis for the observed overall enrichment of deletion expression in ticks, although less supported by current studies, is that POWV D-RNA expression may be associated with persistent replication in the tick vector. Ticks only take bloodmeals preceding transitions from larval, nymphal, and adult life stages, necessitating persistent infection by tick-borne flaviviruses across lifestages for effective transmission to a new host. TBEV has been shown to establish persistent infection in both ticks [30, 34] and tick cells [34], with little evidence of significant genetic variation at the consensus level between parental virus and virus after up to 120 days persistence in *Ixodes* ticks. While previous work has not evaluated the relationship between D-RNA expression and persistence in tick hosts, persistence of LGTV in human embryonic kidney cell culture has been positively associated with the accumulation of defective particles [23]. Thus, D-RNA expression among POWV populations in ticks may be associated with persistent infection of the tick vector, either as a regulator of persistent infection or a byproduct thereof. Although speculative, this hypothesis could be expressly tested by characterizing D-RNA expression over the course of persistent POWV infection in laboratory-infected ticks.

The primary limitation of our study is the use of POWV sequencing data from prior studies not explicitly designed to detect recombination events. We therefore used ClickSeq, a library preparation method designed for accurate and specific detection of recombination, to validate our findings for a subset of samples with residual RNA available. To ensure a rigorous analysis, we only used data from samples that had a minimum average sequencing depth of 100X; above this threshold, we did not observe a relationship between higher sequencing depth and an improved ability to detect recombination. Further, deletions with only one total count were removed in order to remove potential artifacts generated through either the library preparation process or the Illumina platform. An additional limitation is that our analyses were performed with short-read sequencing data; further work is needed to evaluate whether the deletions we detected are present in full-length virus genomes, whether deletions can co-occur on the same RNA molecule, and whether they may be packaged and released as defective particles.

In all, the data presented here offer foundational insight into recombination and deletion within POWV, with possible relevance to other tick-borne flaviviruses. Future studies are needed to understand the potential role of D-RNAs during infection and transmission in both *in vitro* and *in vivo* models of ticks and mammalian hosts. Through continued study of POWV recombination and deletion generation, we can gain further insights into this important pathogen’s evolution and pathogenesis.

## Materials and Methods

### Datasets for secondary analysis

RNA from POWV-positive(+) tick homogenates and single-passage isolates underwent full POWV genome sequencing previously using a random-primed, metagenomic library construction and Illumina sequencing [29, 46]. Datasets were selected for recombination analysis based on the following criteria: 1) RNA or total nucleic acid (TNA) was collected from tick homogenates (of any lifestage); 2) if RNA or TNA was from a tissue culture passage, the original isolate came directly from a tick host or pool and had naturally infected tick-derived sequence data available; and 3) POWV full genome sequencing resulted in a minimum average depth 100.

### ClickSeq library preparation

TNA from tick homogenates was treated with DNase (ArcticZymes) according to manufacturer’s instructions and ClickSeq libraries were generated as previously described [17, 40]. Briefly: RNA was primed with 1uL 100uM primer consisting of random hexamer and partial Illumina p7 adapter sequence, plus 1uL 1:35 azidoNTP:dNTP mix (ClickSeq Technologies), followed by first-strand synthesis with superscript III enzyme (Invitrogen). Template RNA was removed using RNase H (Invitrogen), first-strand cDNA was bead-purified (Ampure bead info), and the full p5 sequencing adapter with a 12 nucleotide unique molecular identifier was added to the first-strand cDNA via Click reaction. Click reaction proceeded at room temperature for two hours, with 2uL 5uM p5 oligo, “ClickMix” (ClickSeq Technologies), and ClickCatalyst (ClickSeq Technologies) prepped with 10mM vitamin C (Biotechne). “Clicked” cDNA was PCR amplified for a total of 21-24 cycles using OneTaq (New England Biolabs), a universal p5 primer, and an i7 indexing primer. Final libraries were run on a 2% agarose E-gel (Invitrogen) and gel-extracted from the 250-650bp range using the Zymo Gel DNA Recovery kit (Zymo Research). Libraries were quantified by QuantIt picogreen assay (Invitrogen), pooled, and sequenced on a NextSeq2000 (Illumina).

### Bioinformatics and recombination analysis

Adapters were removed from reads by Illumina’s BaseSpace software. Raw fastq files were processed using fastp version 0.23.2 [47] to: 1) ensure complete adapter removal; 2) quality filter, deduplicate, filter reads less than 50 bases in length; and 3) merge overlapping reads and write non-overlapping reads to separate read1 and read2 files. For ClickSeq libraries, fastp was additionally utilized to trim and append UMI sequences to read names. Because paired-end sequencing was performed and recombination mapping tools were designed for single-end data, read names in overlapping read, non-overlapping read 1, and non-overlapping read 2 files were tagged with “/3”,”/1”, and “/2”, respectively, and all read files were concatenated into one fastq file for analysis. Reads from tick samples were first mapped to the *Ixodes scapularis* genome using hisat2 v 2.2.1 [48] to remove tick reads. Then, for each sample, a sample-specific POWV consensus sequence was generated by mapping POWV reads to reference sequence HM440559.1 and using pilon [49] (unknown version) to generate a reference-based consensus sequence. POWV reads from each sample were aligned to the sample-specific consensus sequence, and recombination events were identified using ViReMa version 0.25 [50]. Deletions were extracted from resulting BED files, and recombination breakpoint junctions were indexed to the HM440559.1 genome.

### Protein modeling

The amino acid sequence of the POWV strain NFS001 (Genbank HM440559.1) nonstructural protein 5 was submitted to Expasy/SWISS-MODEL [51] to identify closely-matching amino acid sequences available in the Protein Data Bank (PDB) using the HHblits tool [52]. The structure for the Japanese encephalitis virus (JEV) ns5 was identified as a template (4K6M.1.A)[38], with 57% sequence identity and 99% sequence coverage. A model was subsequently built in ProMod3 v3.2.1 [53] via SWISS-MODEL with a resulting QMGE score of 0.84 and a QMEANDisCo of 0.80±0.05. The molecular surface for the resulting model (Supplemental) was calculated in Swiss PDB Viewer v4.1 [54] and amino acids colored by associated nucleotide deletion data.

### Statistics

For analyses comparing total and unique deletions counts, deletions with only one read count were removed and raw counts were normalized to count/10^6^ mapped reads. All statistical analyses were performed in RStudio version 2023.6.2.561 [55] using R version 4.3.1 [56]. Normality was assessed using Q-Q plots in ggpubr version 0.6.0 [57]. The relationship between total read count and either Shannon’s diversity index (H) or total deletion expression was assessed by Kendall correlation analysis. Analysis of variance for multiple groups was performed using the Kruskall-Wallis test, and paired sample analysis was performed using the Wilcoxon rank-sum test. For data that did not fit one or more assumptions of the Wilcoxon rank-sum test, a two-sided paired sample sign test was employed using the EnvStats package for R with mu=0 and confidence interval of 95%. Differential gene expression was assessed using DESeq2 v1.40.2 [58]: for this, all deletions were pooled across samples, and only deletions with at least two counts in at least two samples were included. Raw read counts were used for analysis, without normalization to total mapped reads. The program was run using paired data structure and a local regression fit. DESeq2 output was used to perform principal component analysis (PCA) using prcomp() and for hierarchical clustering analysis in Cluster 3.0 [59]; clustering using average linkage is shown here, although all linkage algorithms resulted in highly similar results. Hierarchical clustering results were visualized in TreeView [60].

## DATA AVAILABILITY

All raw sequencing data will be made available in the Sequence Read Archive (SRA) under BioProject PRJNA1011342.

## ACKNOWLEDGMENTS

This study was supported in part by the Emory Integrated Genomics Core (EIGC) (RRID:SCR_023529), which is subsidized by the Emory University School of Medicine and is one of the Emory Integrated Core Facilities. Additional support was provided by the Georgia Clinical & Translational Science Alliance of the National Institutes of Health under Award Number UL1TR002378. The content is solely the responsibility of the authors and does not necessarily reflect the official views of the National Institutes of Health.

